# Duck slurry as a matrix for avian influenza virus detection and genetic characterization in domestic flocks

**DOI:** 10.1101/2025.09.18.677031

**Authors:** Clément Castille, Laura Lebouteiller, Mathilda Walch, Guillaume Croville, Jean-Luc Guérin, Sébastien Mathieu Soubies

## Abstract

Avian influenza viruses (AIVs) circulate widely in domestic and wild birds and continue to pose major risks to animal and public health, in particular those of high pathogenicity (HP). Ducks play a key role in AIV ecology due to frequent subclinical infections and high viral shedding. Efficient environmental surveillance approaches are needed to complement costly individual testing, especially in the context of large-scale anti-H5 vaccination as implemented in France. We evaluated duck slurry, the wastewater produced in duck fattening units, as a matrix for AIV detection and optimized molecular protocols for sensitive molecular detection and genetic characterization. Slurry samples (n = 172) were collected from 37 duck farms in southwestern France between May 2024 and March 2025. Among several extraction strategies, RNA extraction from the solid fraction using TRIzol LS followed by magnetic bead purification yielded the highest sensitivity. Using this optimized protocol, 82% of samples tested positive for AIV RNA, with 30% of positives being H5-positive; no H7 viruses were detected. Whole-segment RT-PCR and sequencing were successful for shorter genomic segments, enabling subtype determination, although virus isolation consistently failed. These findings demonstrate that slurry is a promising matrix for AIV detection, providing valuable molecular data. Slurry-based surveillance may therefore serve as an effective complement to individual testing and improve early warning capacities for influenza surveillance in a One Health perspective.

**Importance:** In France, current surveillance of high pathogenicity avian influenza in ducks relies mainly on individual swabs, which may miss subclinical or transient infections, particularly with low-pathogenicity strains. By targeting slurry, a wastewater generated in duck fattening units, we developed an affordable and practical approach for environmental surveillance. Slurry samples captured high levels of avian influenza RNA and allowed partial genomic characterization despite failed virus isolation. This strategy is directly relevant to monitoring avian influenza viruses dynamics under current anti-H5 vaccination and can be extended to other livestock systems. Duck slurry-based surveillance thus represents an efficient One Health tool to strengthen preparedness against avian influenza outbreaks.

## Introduction

Avian influenza viruses (AIVs) are enveloped, negative-sense RNA viruses belonging to the *Orthomyxoviridae* family. Their genome consists of eight segments and evolves through both mutation and reassortment. AIVs are subtyped depending on the combination of hemagglutinin (HA) and neuraminidase (NA), the two major surface glycoproteins of the viral envelope. Nineteen HA subtypes and 11 NA subtypes have been described so far, including the recently described H19 subtype (1, 2). All of these subtypes except H17, H18, N10 and N11 circulate among aquatic birds.

AIVs are additionally classified into pathotypes, namely low (LP) and high pathogenicity (HP) depending on the sequence of HA (3). LPAIVs typically replicate in the digestive tract of *Anatidae* without clinical expression (4). In contrast, HPAIVs display systemic replication in ducks with high respiratory, digestive (5) and feather (6) shedding along with variable clinical symptoms depending on viral genotype (7), species (8), age (9, 10) and vaccinal status of birds. Pre-symptomatic shedding in ducks may explain their role as amplifiers of viral outbreaks in high-density poultry areas (11).

Since their emergence, H5Nx HPAIVs of the Goose/Guangdong/96 lineage, particularly those belonging to clade 2.3.4.4b, have caused hundreds of millions of deaths in both wild and domestic birds, infected various mammalian species (including cats and cattle) and led to human cases (12, 13). Extensive reassortment with LPAIVs present in wild birds has resulted in a wide range of genotypes with diverse segment constellations (14).

France is the leading European producer of farmed ducks, that are in particular used for fatty liver production. Ducklings intended for this purpose typically spend their first 80 days of life in farms with outdoor access starting around 40 days of life. Afterwards, they are transferred into indoor fattening units, with a capacity of few hundred individuals each, for a final period of about 15 days. These units, typically equipped with slatted floors, generate a liquid waste - referred to as “slurry” - from faeces and feed residues washed away during cleaning. Slurry is stored on-site and can then be used either as fertilizer or for biogas production.

The high number of duck farms likely explains the numerous HPAIVs outbreaks in poultry farms that France experienced during the recent years. As a result, a nationwide anti-H5 vaccination program for farmed ducks was initiated in autumn 2023. This vaccination scheme is accompanied by thorough but costly virological surveillance to avoid silent circulation of HPAIVs in vaccinated flocks.

Alongside individual testing approaches, environmental virus detection approaches have recently gained momentum, particularly inspired by the success of wastewater-based surveillance of SARS-CoV-2. (15).

The aim of this study was to investigate the interest of duck slurry, a wastewater produced by ducks in fattening units, as an environmental matrix for detecting AIVs infecting ducks to get a better understanding of AIVs circulation in those animals.

## Material and methods

### Samples and collection strategy

Slurry samples originated from thirty-seven duck fattening units situated within 50 km of a methanation plant (Méthalandes, France) in southwestern France. Slurry was serially sampled between May 2024 and March 2025. Slurry from each fattening unit was temporarily stored on-site in slurry pits before being pumped into tanker trucks and transported to the methanation plant. Each truck collected and transported slurry from a single farm at a time. Samples used for analyses were collected before unloading each tanker truck into the plant system, using a manual valve at the back of the tank. Slurry samples (500 mL), collected with a minimal frequency of one collection per month, were kept at 4°C, for a maximal duration of 15 days until transport and processing at the laboratory.

### Optimization of sample treatment and RNA extraction

Aliquots of 1 mL of raw slurry samples were centrifuged at 11,000 g for 1 hour, 4 °C. Supernatant were transferred into new tubes and 5 sample groups were processed (Figure 1A): 1) 125 µL of supernatant underwent RNA extraction using MAGFAST-768 RNA extraction kits (IDVet, Grabels, France) following the manufacturer’s instructions (Figure 1A, “Sup Beads”), 2) 250 µL of supernatant was diluted into 750 µL of TRIzol LS (ThermoFisher) and was treated according to the manufacturer’s instructions until phase separation, when the aqueous phase was either subjected to magnetic bead-based RNA extraction as described above (120 µL aqueous phase) (Figure 1A, “Sup Trizol Beads”) or 3) subjected to RNA precipitation as described below(Figure 1A, “Sup Trizol PP”). Pelleted solids were resuspended into 750 µL of TRIzol LS and incubated for 5 minutes at room temperature. Two hundred microlitres of chloroform were added, tubes were briefly vortexed and incubated for 3 minutes at room temperature, followed by centrifugation for 15 minutes at 12,000 g, 4 °C. The aqueous phase was transferred into new tubes and then treated by either 4) magnetic bead-based RNA extraction (Figure 1A, “Pellet Trizol Beads”) as described above (120 µL aqueous phase) or 5) subjected to RNA precipitation Figure 1A, “Pellet Trizol PP”) as described below.

**Figure 1.**
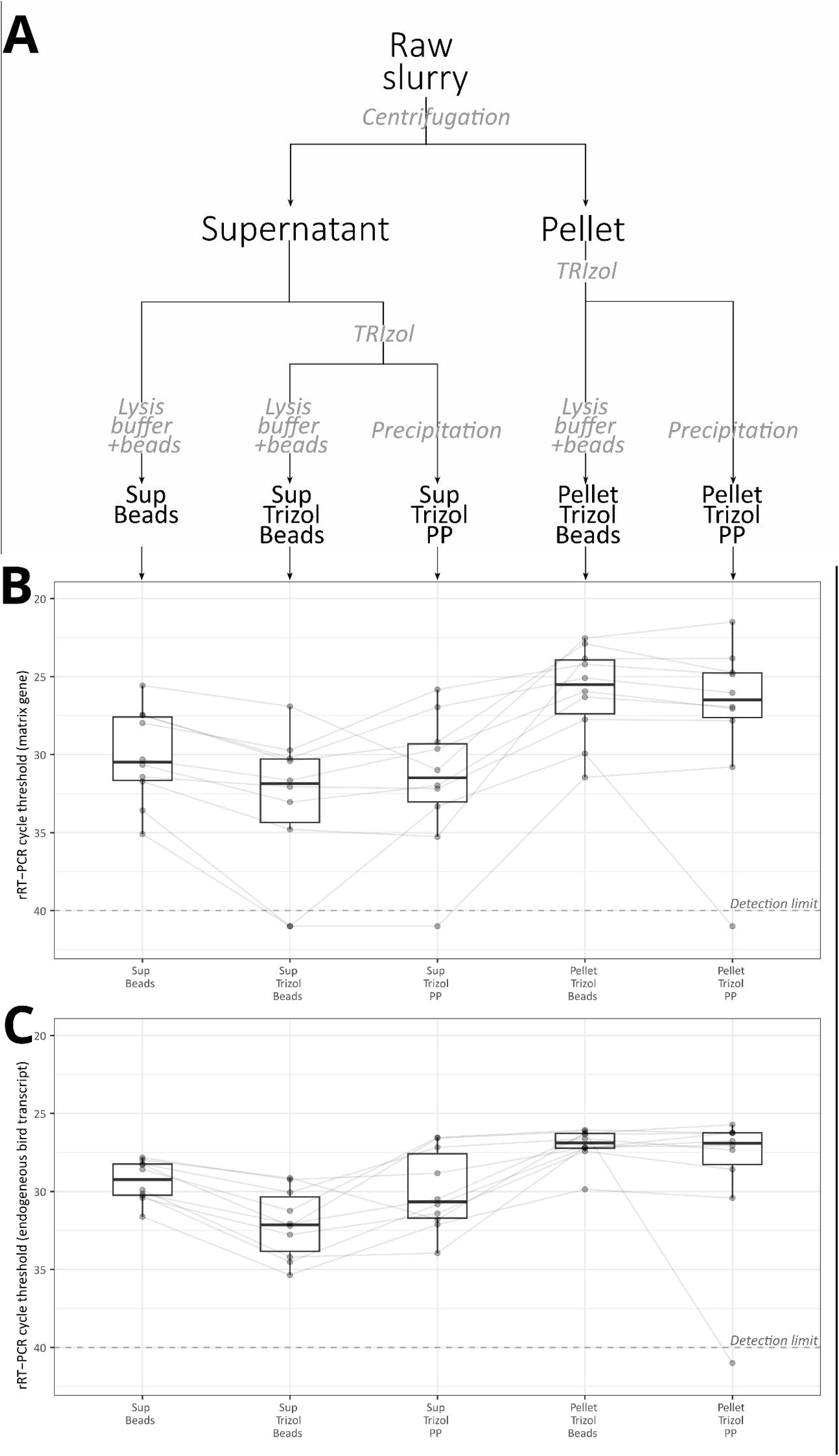
Comparison of influenza RNA (M gene, A) and bird endogenous RNA (B) detection by rRT-PCR using the indicated protocols. Sup: supernatant; PP: precipitation. Boxplots present median, interquartile range and extreme values. Points represent individual slurry samples and lines link values from one slurry sample.

### RNA precipitation

Five hundred microlitres of isopropanol were added to the aqueous phase. Tubes were incubated for 10 minutes at room temperature and centrifuged at 12,000 g for 10 minutes, 4 °C. Supernatants were transferred and pellets were resuspended into 1 mL of 75% ethanol, followed by brief vortexing and centrifugation for 5 minutes at 7500 g, 4 °C. Supernatant were removed and pellets were dried at room temperature until complete evaporation of any residual ethanol. Pellets were then resuspended into 50 µL of ultrapure water.

### Real-time RT-PCR (rRT-PCR)

Real-time RT-PCR (rRT-PCR) was carried out on all extracted RNA using a commercial kit simultaneously detecting the matrix gene as well as an avian reference transcript (reference IDFLUATRI-100 Innovative Diagnostics SAS, Grabels, France). Positive samples were further analysed by rRT-PCR using the IDFLUH5H7-kit (Innovative Diagnostics SAS, Grabels, France) to detect H5 or H7 AIV. All rRT-PCR were performed on a LightCycler96 thermocycler (Roche SAS, Boulogne-Billancourt, France) following the respective kit cycling instructions.

### Full segments RT-PCR and sequencing

Each of the viral segments was amplified by 2-step reverse transcription-PCR (RT-PCR). For reverse transcription, 10 µL of RNA was treated with the RevertAid™ First Strand cDNA Synthesis Kit (ThermoScientific, reference 10387979) using the Uni12 primer (16) according to the manufacturer’s protocol. The obtained cDNAs were used for full-segment amplification using Phusion hot start flex 2x master mix kit (New England Biolabs, reference M0536S) and a set of gene specific primers (16). Eight distinct PCR mixes, corresponding to the 8 genes, were prepared and contained 12.5 µL 2X buffer,7.5µL RNase free water, 2.5 µL of 10 µM primers and 2.5 µL cDNA. The PCR program consisted of an initial 30 s step at 98°C followed by 35 cycles with the following conditions: 98°C for 10 s, 61°C for 30 s, and 72°C for 2 min 20 s. The program ended with 1 step at 72°C for 10 min. PCR products were then excised after agarose gel electrophoresis and sent for “ONT Lite Clonal Amplicon Sequencing” to Eurofins genomics (Ebersberg, Germany).

### Alignment to reference sequences

FASTQ files obtained after Nanopore sequencing were aligned to a set of reference AIV genomes using Geneious Prime 2024.0.7.

### Virus isolation

For virus isolation, 1 mL aliquots of slurry were centrifuged for 1 h at 11 000 g, 4 °C. Three hundred microlitres were collected and mixed with 300 µL of Penicillin/streptomycin (10.000 µg/mL, ThermoFisher), 35 µL of 10% Bovine Serum Albumin (m/V in PBS) and 70 µL of amphotericine B (250 µg/mL, ThermoFisher). Two hundred microlitres of this solution were then inoculated into the chorioallantoic sac (CAS) of 9 to 11 day-old embryonated chicken eggs (2 to 3 eggs per sample). Inoculated eggs were candled daily. After 72 h or until embryo mortality was observed, allantoic fluids were collected and tested by rRT-PCR (M detection) as described above.

## Results

Based on previous data from the laboratory as well as the literature, detection of avian influenza viruses (AIV) RNA by real-time RT-PCR (rRT-PCR) from the supernatant of centrifuged slurry seemed achievable. Initial attempts were therefore performed on slurry samples without viral spiking. This first method, hereafter referred to as the “reference protocol”, involved magnetic bead-based purification of the slurry supernatant after treatment with lysis buffer.

Initial positive results for M gene detection (Figure 1B, column 1 “sup beads”, median Cq: 30,12) demonstrated the feasibility of using naturally contaminated slurry as a matrix for optimizing detection protocols in the absence of viral spiking. Those same samples were then used to compare several extraction strategies (Figure 1A). TRIzol LS - based RNA extraction of slurry supernatant followed by isopropanol precipitation resulted in Cq values comparable to those obtained with the reference protocol (Figure 1B, column 3 “sup trizol pp”, median Cq: 30.59). In contrast, RNA extracted from slurry solid fraction (pellet) resuspended in TRIzol LS and subjected to isopropanol precipitation yielded significantly lower Cq values (Figure 1B, column 5 “pellet trizol pp”, median Cq: 25.95). To circumvent the lengthy steps associated with isopropanol precipitation, a variation of the TRIzol LS protocol described above was tested, both on slurry supernatant and pellet. In this modified protocol, the aqueous phase obtained after chloroform extraction was subjected to magnetic bead-based RNA purification: the resulting Cq values were comparable to those obtained after isopropanol RNA precipitation (Figure 1B, column 2 and 4 “sup trizol beads “ and “pellet trizol beads”, median Cq of 31.1 and 25.99, respectively). The “pellet trizol beads” protocol, combining high analytical sensitivity with ease of implementation, was considered as the “optimized protocol” and used in subsequent experiments.

Real-time RT-PCR analysis of the endogenous RNA detected using the commercial kit showed a trend consistent with that observed for AIV RNA, albeit with a milder decrease in Cq between the reference protocol and the optimized protocol, (Figure 1C, column 1 versus 4).

A total of 172 slurry samples collected from 37 duck farms between May 2024 and March 2025 were subsequently analysed using the optimized protocol. Among the 37 farms from which slurry was analysed, 23 farms provided at least 3 samples during the study period. Overall, 142 samples (83%) tested positive for AIV RNA (Figure 2A). Among those positive samples, 43 tested positive for H5, corresponding to 30% of AIV positive samples and 25% of all analysed samples. No sample tested positive for H7 AIV. A significant lower median Ct was observed for M+H5+ samples (median value: 28.22) when compared with M+H5-samples (median value: 31.55, p=2.1e-5 using Wilcoxon test, Figure 2A).

**Figure 2.**
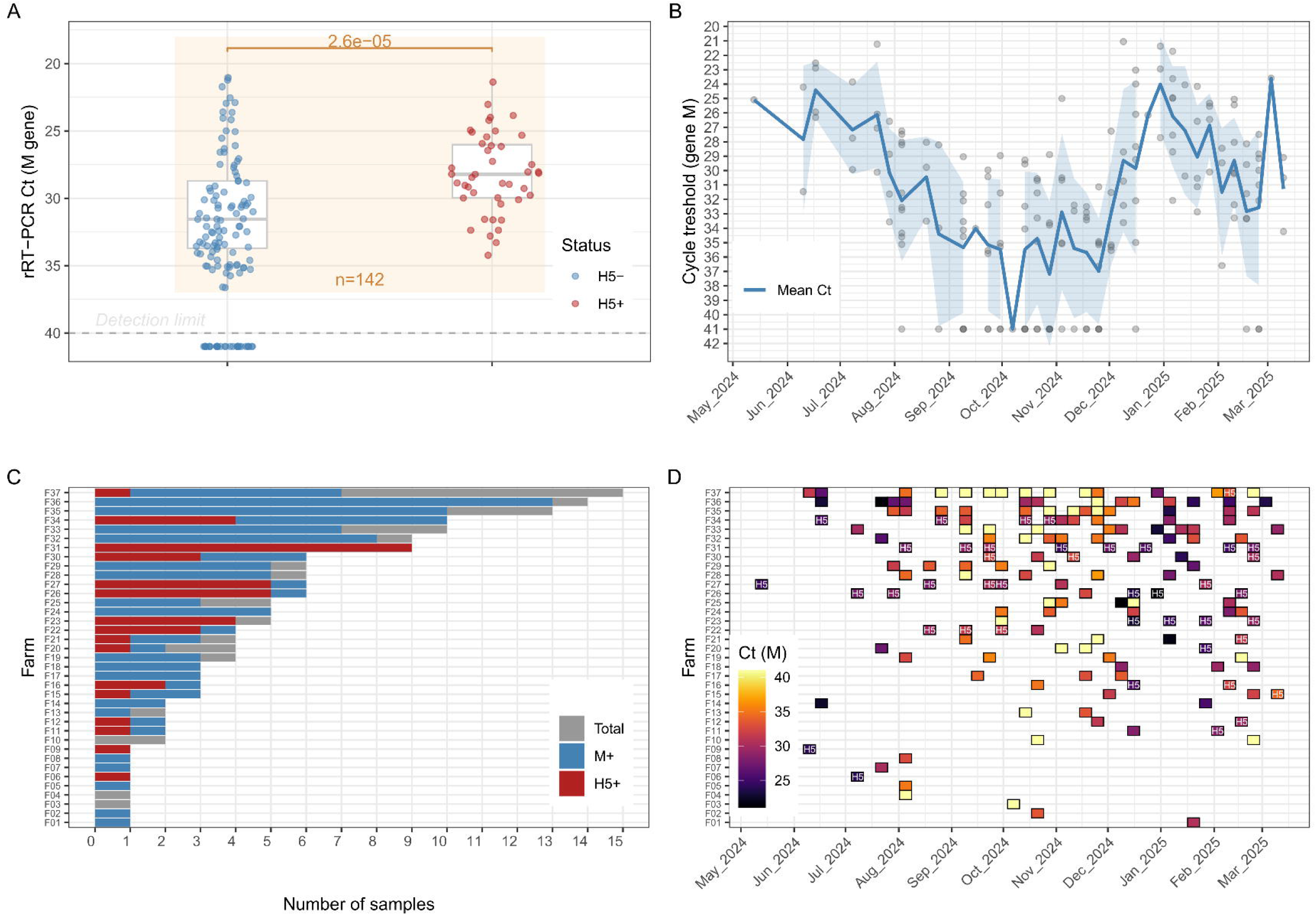
rRT-PCR detection of avian influenza virus (AIV) in slurry samples from 37 farms between May 2024 and March 2025. A: Distribution of M gene Ct values in analyzed samples (n=172); boxplots show distributions for M positive, H5-negative samples (left) and M positive, H5-positive samples (right); the p-value corresponds to a two-tailed Wilcoxon test. B: Temporal variations of mean M gene Ct values (line). The shaded area represents the standard deviation and points represent individual samples. C: Number of samples provided by individual farms, showing total samples (grey), M-positive samples (blue), and H5-positive samples (red). D: Temporal patterns of AIV detection in individual farms; the colour scale indicates M gene Ct values and H5 positive samples are indicated. For all panels, negative samples were empirically assigned a Ct value of 41.

Temporal variations in Ct values were observed (Figure 2B), with mean values around 25-30 observed in June-July 2024 followed by an increase above 30, between August and December 2024, consistent with lower genomic viral loads. Mean Ct values decreased again in January - February 2025. Farm-to-farm variations were observed in the number of collected samples (Figure 2C). Thirty-four out of the 37 studied farms provided at least one positive slurry sample, and 16 farms provided at least one H5-positive sample. Among farms with at least five analyzed samples (n = 15), distinct patterns of AIV detection were observed. Six farms (F24, F26, F27, F30, F31and F34) were positive for AIV in all collected samples. Among those, four farms (F26, F27, F30 and F34) yielded both M+,H5- and M+,H5+ samples (Figure 2C). Longitudinal analysis of individual farms with at least 5 samples further revealed that for 8 out 15 farms, at least 2 series of positive samples separated by at least one negative sample were observed (Figure 2D).

Whole-genome RT-PCR amplification was attempted on a subset of samples with M gene Cq values below an empirical cut-off of 25, using RNA extracted with the optimized method. In several cases, RT-PCR yielded visible amplicons on gel electrophoresis for segments 4 to 8 while amplification failed for segments 1 to 3 (Figure 3A). Shorter segments generally yielded higher amplification efficiency, although amplicons for segments 4 and 6 were occasionally undetectable. Visible PCR products were excised, gel-purified and subjected to Nanopore sequencing. Consensus sequences were obtained after alignment with a set of reference influenza genomes, in particular for segments 4 and 6, which encode HA and NA, respectively, for subtype determination. In the representative result shown (Figure 3B), sequencing depths of several hundred reads were obtained for segments 4 to 8, enabling robust consensus sequence generation. Virus isolation was attempted on samples whose Cq values for M gene were lower than an empirical cut-off of 32. Out of 39 sampled tested during the study period, none yielded AIV isolates.

**Figure 3.**
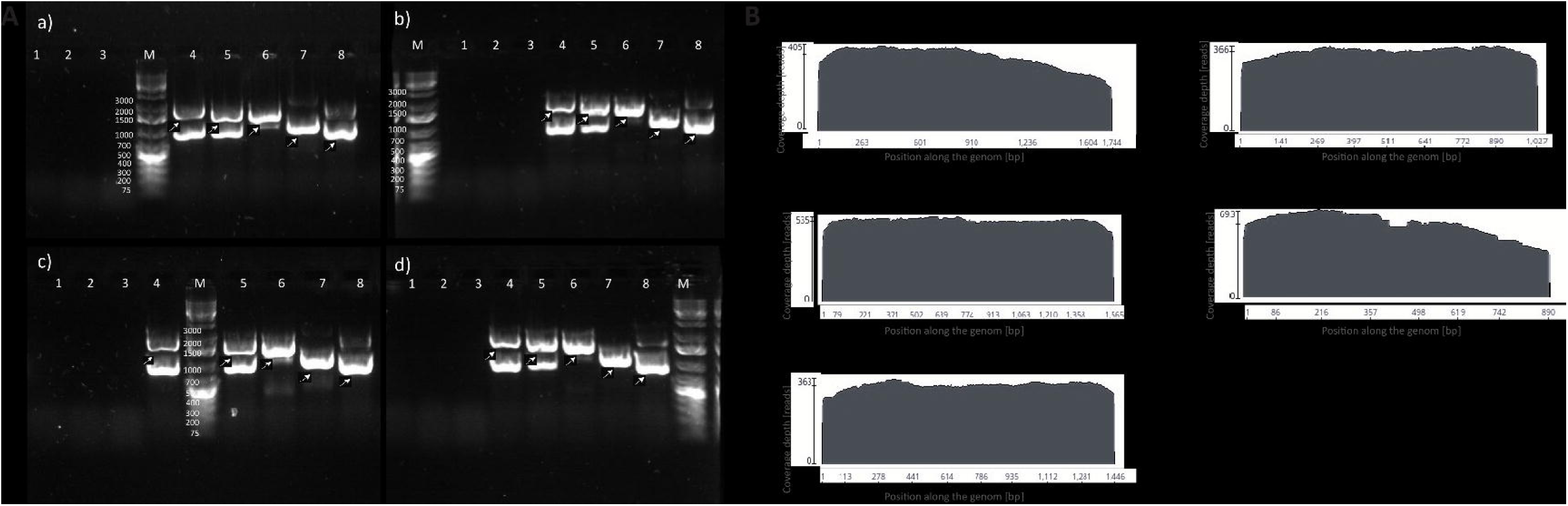
Genome RT-PCR amplification (A) and sequencing (B) of avian influenza viruses from slurry samples. A: representative agarose gel electrophoresis of sets of PCR amplicons from four slurry samples (a) to d)). 1-8: viral segment number; M: molecular ladder. B: example of sequencing coverage plots presenting read number per segment position from segments 4 to 8. Reads were mapped to reference A/Mallard/Sweden/SVA250129SZ0398/JM026682/OT/2024.

## Discussion

In this work, we optimized the detection of avian influenza viruses (AIVs) in duck slurry to enable viral surveillance and genetic characterization. Molecular detection using the slurry supernatant had previously been performed by our laboratory (unpublished results) and others (17). Based on other reports on influenza detection in soil (18) and in wastewater (19), we found that RNA extraction from the solid fraction of slurry using TRIzol LS resulted in enhanced detection sensitivity by rRT-PCR. Combining TRIzol LS extraction with magnetic bead-based RNA purification further improved sample workflow. The reasons for preferential detection of influenza RNA in the solid fraction remains unclear, although influenza viral particles have been reported to adsorb on particulate matters (20). Interestingly, the endogenous duck RNA detected using the rRT-PCR kit was also more abundant in the solid fraction.

Longitudinal analysis of slurry collected in the methanation plant revealed a surprisingly high frequency of AIV-positive samples, including approximately one-third that were H5-positive. Although each truck collected slurry from a single farm at a time before unloading into the plant system, the absence of tank cleaning between successive collections may have led to spillover contamination, thereby artificially inflating the observed positivity rate. For instance, assuming a hypothetical residual volume of positive slurry of 2 % of the tank capacity, the next sample collected might yield a false positive result with a Cq value approximately 5 cycles higher. Careful analysis of truck collection sequences may provide more accurate insights into the true positivity rate of the samples, which is beyond the scope of this study.

The high detection rate of H5 sequences in the collected samples raises the question of the pathotype of the detected viruses. Although formal classification would require sequencing of the HA cleavage site, which is beyond the scope of this work, several lines of evidence suggest that the detected viruses were most likely of low pathogenicity: i) only four H5 HPAIV outbreaks were reported in isolated duck farms in southwestern France during the study period, ii) the thorough official surveillance scheme enforced following the implementation of anti-H5 vaccination in ducks did not report H5 HPAIV positive detection in the area surrounding the methanation plant during the period of our study, iii) partial HA sequencing from one H5-positive sample revealed a monobasic cleavage site consistent with a LPAIV (data not shown). The absence of H5-positive reports in the region—despite numerous positive detections in our samples—may seem paradoxical. A likely explanation is that individual surveillance in vaccinated duck flocks relies primarily on oropharyngeal or tracheal swabs. These samples are excellent indicators of respiratory shedding of H5 HPAIVs, particularly those of clade 2.3.4.4b, which are the main target of current surveillance programs in France. However, in the case of LPAIV infection in ducks, respiratory shedding is only transient (21), and such infections may be missed using oropharyngeal swabs alone. Considering that H5 LPAIV circulation in domestic ducks may be a prelude to the emergence of HPAIVs, a scenario already experienced in France (22), slurry-based detection may serve as a low-cost, effective complement to individual surveillance.

Using slurry as a matrix to detect AIVs suffers from several limitations. First, it is applicable only to farm systems that produce liquid waste, such as ducks in fattening units, thereby excluding earlier production stages when birds are not housed under such systems. Second, as mentioned above, cross-contamination between successive samples or over time on a given farm may occur. Third, in our hands, virus isolation and amplification, which would render complete genome sequencing possible, has consistently failed. The failure to isolate virus - in contrast with the findings of Schmitz et al. - may be explained by several factors: i) the likely LPAIV nature of the detected viruses, which may results in lower levels of viral shedding in the faeces compared with HPAIVs, ii) the acidic pH of some slurry samples (pH <6, data not shown), which may rapidly inactivate virions, iii) warm ambient temperatures during parts of the study period (spring to autumn 2024), which may have accelerated viral degradation, and iv) delays of up to 20 days between sample collection and virus isolation attempts. Despite these limitations, slurry-based AIV detection appears to be a promising surveillance tool. It can provide valuable genetic information and potentially track the emergence of HPAIVs from LPAIVs progenitors. Future work will aim to generate more detailed temporal data on AIV dynamics at the farm level, along with deeper genetic characterization of the detected viruses. More broadly, such approaches could be extended to other livestock systems and pathogens, contributing to integrated One Health surveillance strategies.

## Acknowledgement

The authors wish to thank Mr. Xavier Labat, owner of the methanation plant, and Méthalandes technical staff who helped for slurry sample collection.

## Funding

This work was supported by funding from from the French government, managed by the National Research Agency (ANR), under the France 2030 program OBEPINE+ (reference ANR-24-MIEM-0004), “Chaire de professor junior “ held by Sebastien Mathieu Soubies, as well as by the French “Région Nouvelle-Aquitaine” (project EMERG) and the SISP&EaU project (reference ANRS-23-PEPR-MIE-0003), funded under the PEPR-MIE program and managed by ANRS-MIE (French Agency for Research on AIDS and Viral Hepatitis – Emerging Infectious Diseases).

## Data availablity

The raw data supporting the findings of this study are available from the corresponding author upon reasonable request.

## Author contribution

Conceptualization: SMS, JLG. Methodology: CC, LL, MW, GC, SMS. Investigation (sample collection, laboratory work, sequencing): CC, LL, MW. Data curation & Formal analysis: CC, LL, SMS, MW, GC. Writing – original draft: CC, LL, SMS. Writing – review & editing: CC, LL, SMS, GC, JLG, MW. Supervision: SMS. Funding acquisition: SMS, JLG.

## Conflict of interest

The authors declare no conflict of interest.

## Ethics statement

Slurry samples were collected at the methanation plant with the consent of the owner. No live animals were sampled for this study. Virus isolation was performed by inoculation of embryonated chicken eggs, a standard method that does not require institutional ethics approval under European and French regulations for laboratory diagnostics.

## Notes

### Competing Interest Statement

The authors have declared no competing interest.

### Summary of Updates

In this revision, Figure 2 has been expanded to provide a more detailed description of the temporal series of samples collected between April 2024 and March 2025. A statistical analysis has been added to the M gene Ct distribution previously shown (now Figure 2A). Temporal variations in viral genomic loads are now presented both at the global level (Figure 2B) and at the individual farm level (Figure 2D). In addition, cumulative numbers of H5 and non-H5 positive detections are displayed for each farm (Figure 2C). The corresponding Results section has been updated accordingly.

